# FoldaVirus, a knowledge-based icosahedral capsid builder using AlphaFold

**DOI:** 10.64898/2026.03.27.714795

**Authors:** Oscar Rojas Labra, David S. Montoya-Munoz, Nelly Santoyo-Rivera, Jeffrey McDonald, Daniel Montiel-Garcia, David A. Case, Vijay S. Reddy

**Author notes:** Correspondence to: Vijay S. Reddy, Ph.D., The Hormel Institute, University of Minnesota, 801 16^th^ Avenue NE, Austin, MN 55912, USA.

## Abstract

Coat protein (CP) tertiary structures and their capsid organization of spherical viruses are highly conserved within each virus family. While AlphaFold successfully predicts the tertiary structures of individual CPs, their association to form proper quaternary assemblies cannot be easily accomplished. Here, we report a generalized methodology and associated web-based utility (https://foldavirus.org) that combines AlphaFold predictions of CPs with the knowledge on corresponding icosahedral architectures (e.g., *T*=1, 3, 4…) based on the known structures from the same virus family to generate associated capsids. The resulting assemblies are subjected to Amber energy minimization to relieve any steric clashes at the inter-subunit interfaces. Significantly, the capsid models are validated by calculating robust Mahalanobis distance using the residue annotations categorized as interface, core and surface amino acids with respect to those observed in the experimentally determined analogous structures. Given the amino acid sequence of CP(s), we successfully generated capsids up to *T*=9 icosahedral symmetry, including those of Picornaviruses that display pseudo-*T*=3 symmetry comprising different CPs. As the number of currently available CP sequences are 3-4 orders of magnitude larger than the experimentally determined 3D-structures, this approach bridges the huge gap that exists between the corresponding sequence and structure space.

## Introduction

Virus capsids play multiple roles in the viral lifecycles that include receptor-mediated cell entry, endosomal escape and delivery of viral genomes to the cytoplasm (e.g., ssRNA viruses) or trafficking of nucleocapsids to the nucleus (e.g., dsDNA viruses), viral assembly and genome packaging in producing progeny virions^1-4^. Therefore, structural knowledge of viral capsids provides valuable information in combating viral infections by interfering with their “programmed” virus-host interactions and other key events in the virus lifecycle. Additionally, the structural information can be also used for rational vaccine design^5^ and to identify broadly neutralizing antibodies across a group of similar viruses ^6-9^. However, there exists a large gap between the available viral coat protein (CP) sequences and the known virus structures as characterizing them experimentally is time consuming, expensive, and in some cases not even possible.

In maintaining Virus Particle ExploreR database (VIPERdb; https://viperdb.org) ^10, 11^, a repository of experimentally determined icosahedral virus capsid structures and knowledge base of structure-derived properties, we and others have observed that CP structures and their capsid architectures are highly conserved within each virus family ^12, 13^. For example, while the members of Parvoviridae family always form *T*=1 capsids, the viruses from Picornaviridae family assemble into particles exhibiting pseudo-*T*=3 icosahedral symmetry. The latter *T*-numbers denote *Triangulation* (*T*) number, which is defined as *T*=*h*^*2*^*+hk+k*^*2*^, where *h* and *k* are integers and the indices of a hexagonal lattice ^14, 15^. In other words, not surprisingly, the CP sequence determines the type of capsids they are likely to form. At the time of this writing, there are ∼1,700 viral capsid structures from 92 different virus families and 203 genera available in VIPERdb^10^. In addition to documenting various structure-derived metadata, in VIPERdb, the viruses are grouped according to their taxonomy, genome type and capsid architectures, characterized by their *T*-numbers. Notably, the structure-derived metadata include capsid diameters, net surface charge, buried surface area (BSA) based association energies of the unique subunit interfaces and the annotations of CP amino acids as surface, core, and interface residues^16, 17^.

While AlphaFold2 (AF2)^18, 19^ or AF3^20^ are known to predict the tertiary structures of individual CPs accurately and sometimes their multimer organization, it is currently not possible to build complete models for even simple (*T*=1) capsids composed of 60 copies of a CP, due to GPU memory limitations^21^. In addition, identifying the relevant sub-assemblies from the various degenerate configurations is challenging (e.g., closed hexamer of protomers vs. pentamer + a protomer). While there have been published reports of building exclusively *T*=1 capsids of adeno associated viruses (AAVs) based on the CP sequence^22^, there are no other known reports, to the best of our knowledge, describing similar prediction of capsids displaying different *T*-numbered architectures. To address this limitation, we developed a hybrid method that involves obtaining AF model(s) for given CP sequence(s) and assembling their proper icosahedral asymmetric units to generate complete capsids according to the known quasi-equivalent icosahedral architectures (e.g., *T*=1, 3, 4…) observed in the respective virus families. Furthermore, these capsids are relaxed using Amber energy minimization, thereby relieving any steric clashes that may occur between the neighboring subunits, while assembling the complete capsids. Finally, the capsid models are evaluated based on various structural metrics (e.g., pTM, pLDDT, TM-score) and importantly validated by calculating the robust Mahalanobis distance based on the structure-derived metadata and comparing them with those of the known structures in the same virus family.

We have successfully applied the above method for a number of test cases and evaluated the resulting models against the known structures with considerable success. We report here the details of the methodology and corresponding web-tool (https://foldavirus.org), which is freely available for the scientific community. We believe that this approach provides a way forward to bridge the large gap that exists between the known CP sequences and the limited number of virus/capsid structures determined by the classical structural biology approaches – cryo-electron microscopy (cryo-EM) or X-ray crystallography^10, 23^.

## Results

### Overview of FoldaVirus workflow

Leveraging the observation that the CP tertiary structures and their quaternary organization are highly conserved within each virus family (Supplementary Fig. 1) ^12, 13^, the FoldaVirus pipeline combines the AF predictions of CPs with a knowledge-based approach guided by the known virus structures, their CP sequences, observed quasi-equivalent icosahedral architectures (*T*-numbers) and the associated virus taxonomy to generate relevant capsid assemblies. In the event of observed polymorphism, where a CP is known to form multiple types of capsids (e.g., *T*=1, 3, 4), the user is given a choice to select the capsid of interest to build.

Figure 1 shows the workflow of FoldaVirus methodology. Briefly, the user submitted CP amino acid (a.a.) sequence is searched against a local BLAST library built from the a.a. sequences of CPs of the known and curated capsid structures available at VIPERdb ^10, 11^. This search identifies the types (*T*-numbers) of capsids that the input sequence is likely to form based on the sequence similarity of related structures in VIPERdb. The closely identified structure will be used as the reference (template) structure to build corresponding *T-*numbered capsid for the input sequence (see below). Moreover, we also provide a list of closely related structures from the same virus family, if the user prefers to use a different reference structure. This option is particularly useful that allows the user to build a model of a particular capsid state (e.g., full capsid vs. altered capsid or empty capsid of Picornaviridae; different expansion intermediates of HK97-like capsids etc.).

**Figure 1.**
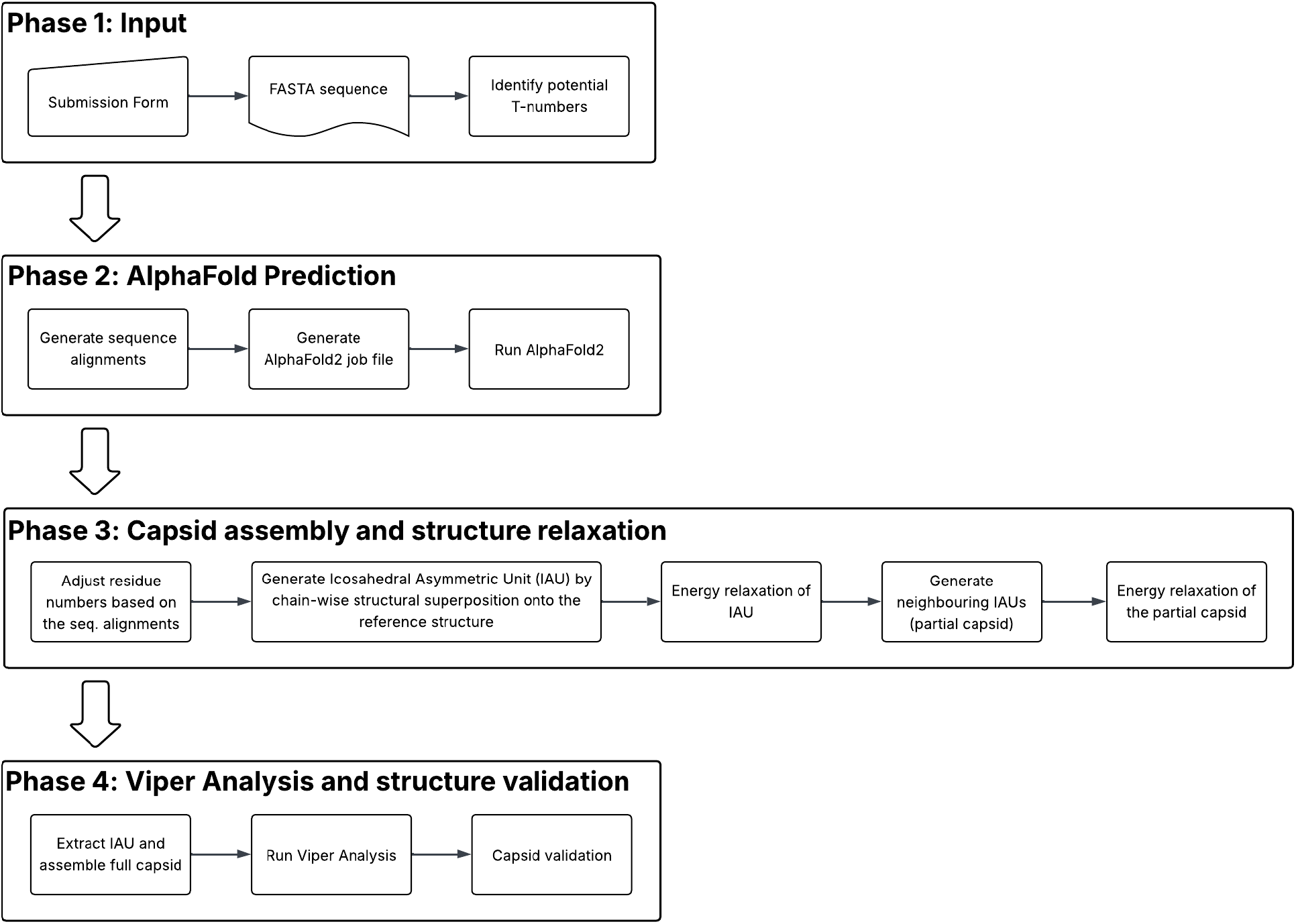
FoldaVirus pipeline. The workflow showing various steps involved in FoldaVirus capsid prediction using AlphaFold.

From the identified hits and the associated sequence alignments, the input sequence is trimmed by removing the likely unstructured (disordered) regions, usually located at the N- and C-termini of the CP structures. Generally, AlphaFold (AF) is known to generate “spaghetti-like” models for the unstructured regions of the sequences. Such unwieldy structures can cause significant steric clashes when generating the complete capsid assemblies, therefore such regions were removed prior to proceeding with the subsequent steps in capsid prediction (Supplementary Fig. 2). The trimmed input a.a. sequence replicated by the number of copies of CPs that occupy an icosahedral asymmetric unit (IAU), as necessitated by the *T*-number of the chosen capsid and submitted to obtain AF predictions. The AF2 docker modules were built on our local HPC cluster at the Hormel Institute and the entire pipeline is run as a batch (SLURM) job. Of note, while we built a similar pipeline that uses AlphaFold3 (AF3), we are unable to release it publicly due to constraints on the terms of use of AF3, even though AF3-pipeline is 3-4 times faster than that using AF2.

The residue numbers of resulting FoldaVirus models are renumbered to match the original amino acid sequence and oriented in the VIPER (Virus Particle ExploreR) convention^24^ by structural superposition, followed by Amber energy minimization of the IAU sub-structure. The above renumbering of residues is necessary to make amends for the trimmed sequences of unstructured regions that were removed previously, as AlphaFold assumes the input sequence always starts at residue number 1. Subsequently, we generate a partial capsid by identifying and including the subunits surrounding the central IAU according to the standard icosahedral symmetry (VIPER convention) (Supplementary Fig. 3), using Oligomer_Generator API from VIPERdb, and again subjected to Amber energy minimization to relax potentially clashing residues at the inter and intra-IAU subunit interfaces. The central IAU is extracted from the ensuing relaxed partial capsid structure and used to generate the full capsid by applying the standard 60-fold icosahedral symmetry matrices. Based on the resulting capsid, various VIPER analyses are performed that include generating contact tables, calculating buried surface areas and the corresponding association energies at the unique subunit interfaces^17^. Furthermore, as a way of validation, the results from VIPER analysis of the FoldaVirus derived capsid models are compared with those of the known capsid structures in the respective virus families available at VIPERdb (see below). All in all, the workflow contains a total of 20 steps, an abridged version of them is shown in Figure 1. It is implemented in a way that a job can be executed up to a specific step (e.g., step #10) or restarted from any step, in case a job was stalled at a certain step for any reason. Such an implementation adds efficiency in executing the workflow. Notably, all the metadata associated with various steps of each prediction are stored in a behind the scenes MySQL database, which are used for running cron jobs in the background to monitor/control the execution of the FoldaVirus workflow.

Each user submitted sequence is given a unique identifier (e.g., fv-#####), using which the results can be accessed from the dynamically generated webpage for each successful submission. In the event, the user wants to keep his/her submission private, a button is provided at the bottom of the page, near the submit button, to keep the submission unlisted, hence cannot be accessed from the “browse available entries” page. However, it can be accessed readily if and when the user shares the unique job identifier or the webpage link with others. The submissions based on the UniProt identifiers are distinctly represented with their UniProt-IDs and the associated *T*-numbers so as to avoid repeat submission of the same sequences. In addition to various structure-based analysis of capsid models that are organized in separate tabs, the results page also contains a molecular graphical display of the modeled capsid structure using the web application of Mol* program ^25^ as well as links to download the coordinates of the models.

Of note, the current implementation of FoldaVirus restricts capsid generation of those with *T*-numbers less than or equal to 9. Moreover, in the event a user accidentally uploads the sequence of a non-capsid forming protein (e.g., Hemoglobin), the knowledge-based system identifies such cases and responds to the effect that no permitted capsids can be built from the uploaded sequence.

### FoldaVirus web interface

The FoldaVirus website can be accessed from the URL, https://foldavirus.org. The web interface is shown in Supplementary Fig. 4. The only inputs required for a FoldaVirus prediction are the a.a. sequence(s) of the CP(s) in FASTA format and the contact information of the user. Alternatively, the user can also simply provide the UniProt-ID of the CP. If the user wants to make changes to the original sequence (e.g., mutations, insertions or deletions), they can modify the sequence in the input window accordingly before submitting the job. The behind-the-scenes knowledge-based system reads in the a.a. sequence and suggests the type of capsid(s) that the input CP sequence is likely to form. Once the user submits the required inputs, the web-interface provides the user with a job identifier, using which the user can access the results page (Fig. 2) once the job is completed successfully. Additionally, the user is informed of the details of the job along with a hyperlink to access the results via the email provided or the status of the job in case the job has failed or aborted. To complete a prediction, it usually takes anywhere between 30 min – 4 hrs depending on the size of the CP and the *T*-number of the capsid being assembled, bigger these numbers, longer it takes. Moreover, based on the identified (potential) *T*-number, the knowledge-based system automatically detects if the capsid contains multiple CPs (e.g., pseudo-*T*=3 capsids of Picornaviridae) and prompts the user to input a.a. sequences for different proteins (e.g., VP1, VP2 and VP3) separately. Once the prediction is successfully completed, the user can view the structural metrics of the prediction and the results of various capsid analysis of the model and download the coordinates of IAU from the results page (Fig. 2).

**Figure 2.**
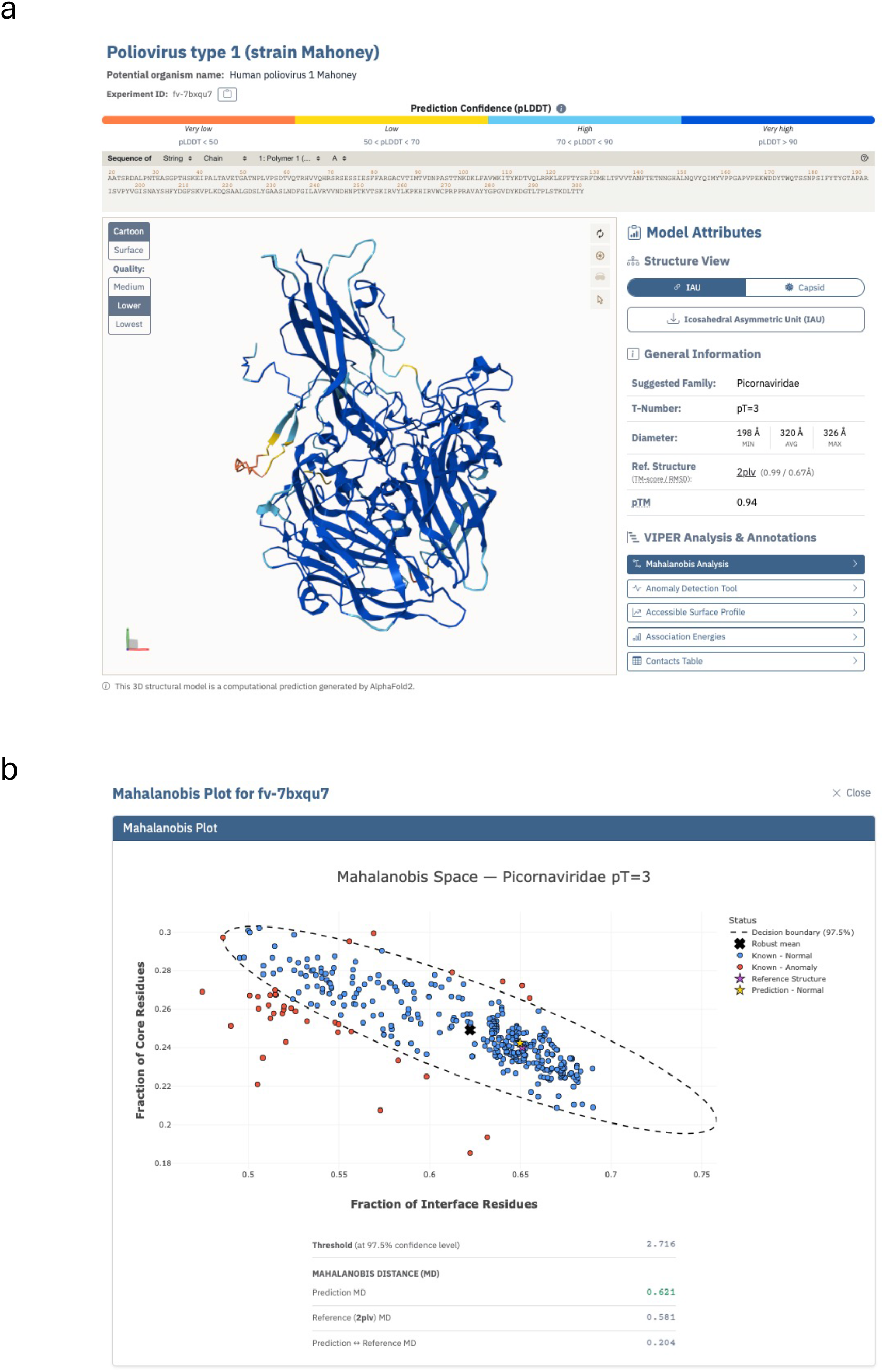
A representative results page of FoldaVirus predicted capsid model. A) Depicted on the left is the predicted model (IAU) shown in the graphical user interface of Mol* web application^25^. Users can toggle between IAU and full capsid representation using the buttons provided in the right panel. Shown on the right are various attributes of the predicted model, links to download the IAU coordinates and the results of various capsid (VIPER) analysis that are organized in separated tabs. B) A Mahalanobis plot, calculated based on the predicted capsid model attributes, showing the agreement of the predicted model in comparison to the known structures in the same virus family. The relevant Mahalanobis distances and thresholds are shown below the plot.

### Limitation of AlphaFold in predicting the oligomers that represent proper icosahedral asymmetric units

While AF2 or AF3 correctly predict the individual CP structures, they are limited in their ability to generate the correct oligomer that represents the proper IAU. This is particularly manifested in the cases of larger quasi-equivalent capsids with *T*-numbers >= 7, sometimes we observed this limitation even in the case of T=3 capsids. In these cases, the resulting oligomers provided by AF cannot be directly used to generate complete capsids by applying the icosahedral symmetry (Fig. 3). We overcame this limitation by structurally superimposing the individual chains onto the corresponding ones in the reference (known) structures, identified by the FoldaVirus pipeline, thereby generating the correct IAU in the VIPER standard orientation^24^. The “rebuilt” IAU is energy minimized independently and in the context of the subunit neighbors (i.e., partial capsid) and used it to generate a complete capsid as described below.

**Figure 3.**
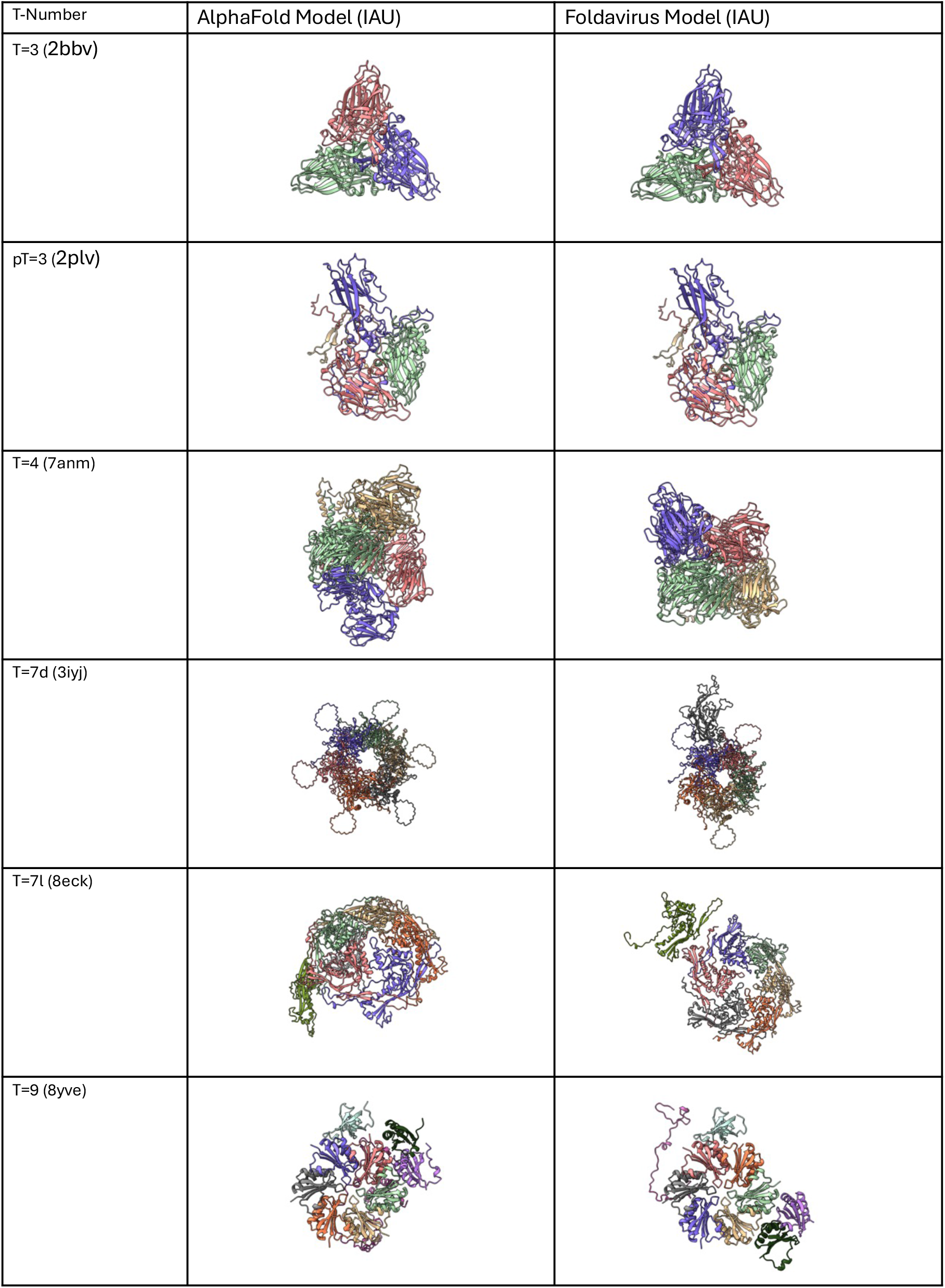
Comparison of the predicted IAUs of representative capsids exhibiting different icosahedral architectures (*T*-numbers) provided by AlphaFold2 with the corresponding IAUs from FoldaVirus. The *T*-numbers of IAUs are shown in column 1 and the corresponding reference PDBs are indicated in parenthesis. The ribbon diagrams of IAUs from AlphaFold2 predictions are shown in column 2, which do not yield complete capsids upon applying the standard icosahedral symmetry operators. However, the adjusted IAUs from FoldaVirus (column 3) can be used to generate proper icosahedral capsids. Of note, the different IAU structures indicated are not drawn to the scale.

### Energy relaxation of FoldaVirus predicted capsid models

Both the IAU and partial capsid models were subjected to two rounds of energy relaxation using the sander program in AmberTools^26^. In the first round, 100 cycles of energy minimization were performed having all the atoms with the exception of Hydrogen atoms restrained (restraint_wt=10.0) with 50 cycles of Steepest Descent, followed by 50 cycles of Conjugate Gradient minimization. In the second round, another 100 cycles of energy relaxation were performed with only the backbone atoms (CA, N, C) being restrained (restraint_wt=5.0). Similar relaxation regimen was applied to the partial capsid consists of the subunit neighbors surrounding the central IAU (Supplementary Fig. 3). The relaxed model of IAU, extracted from the energy minimized partial capsid, is used to generate the complete capsid and for performing subsequent structural (VIPER) analysis.

### Annotation of a.a. residues as interface, core and surface residues

As part of the VIPER analysis of icosahedral capsids, we estimate buried surface areas (BSA) of the CPs at unique subunit interfaces within and surrounding the reference IAU^17^. This involves estimating the solvent accessible surface areas (SASAs) of individual a.a. acid residues in the CPs comprising the IAU, in the context of subunit neighbors. Based on this information, we distinguish whether an amino acid residue is likely to be located at the interface, core or on the inner/outer surface of a CP using the formulae given below. SASA_reference_ refers to SASA of an amino acid in its free (isolated) state, while SASA_bound_ and SASA_unbound_ correspond to SASAs of an amino acid when interacting and not interacting with its subunit neighbors, respectively.

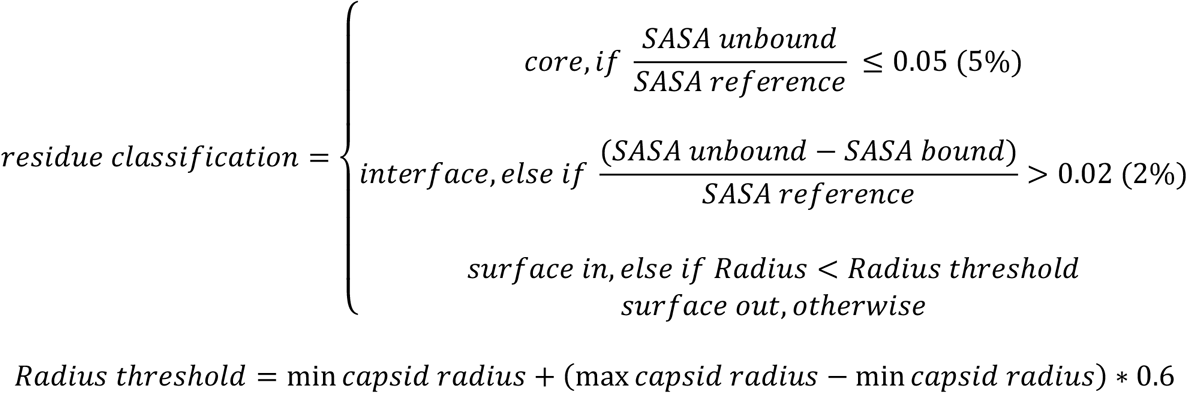

The number of amino acids at different regions (interface, core and surface) were used to calculate Mahalanobis distance to validate the FoldaVirus models as described below.

### Validation of FoldaVirus generated virus capsid models

The validation of FoldaVirus models involves comparing them to known capsid structures from the same virus family and the *T*-number by calculating Mahalanobis distance (MD) using the CP residues that are classified as core or interface residues. The surface residues were not considered in this analysis, as they are dependent on the sum of above two classes. MD is regarded as the “gold standard” for identifying outliers in a complex dataset^27-29^, therefore provides a good metric for validating the FoldaVirus models with respect to the distribution of known structures from the same virus family, as the tertiary structures of CPs and their capsid organization are highly conserved within the same virus family (Supplementary Fig. 1). We used the normalized fractions of core and interface residues relative to the total number of residues in the IAU to calculate MD. The distribution of interface, core and surface residues of CP subunits in the context of an assembled capsid represent a characteristic of particular type of quaternary organization that are generally conserved in each virus family or genus^30^. To calculate the robust distribution of similar structures, we employed Minimum Covariance Determinant (MCD) estimator by minimizing the covariance matrix of the subset of closely matching observations (structures) to identify the outliers. An anomaly threshold is calculated from the chi-squared distribution at 97.5% confidence level. MD below this anomaly threshold indicates that model belongs to the main distribution of structures, hence considered a good model, while MD above the threshold indicates that model could be an outlier. In addition to calculating the conventional MD between the robust mean and the model, we calculated the pairwise MD between the predicted model and the closest (reference) structure that we identified from the set of known structures. The MD between the model and the reference structure is particularly a useful measure, particularly when the reference structure belonging to a different genus is itself classified as an outlier among the known structures from the same virus family. For example, the capsids of Densovirus genus in Parvoviridae family fall outside the main distribution of the majority of capsids that belong to Dependoparvovirus genus (Supplementary Fig. 5).

## Discussion

The underpinnings of workflow design and methodology of FoldaVirus allows the users to seamlessly generate models of icosahedral viral capsids based on CP sequences of virus families for which some structural information on the type of capsid architectures that they are likely to form is known. In addition to generating the capsid models, we have a number of metrics in place to validate them. These include the standard structural metrics like pTM, pLDDT (from AlphaFold) and TM-score with respect to closely matched known structure and the estimated robust Mahalanobis distance (MD) based on the normalized fraction of residues annotated as interface and core residues with respect to the known structures from the same family. Significantly, MD represents the similarity measure of quaternary organization of the predicted capsid model with respect to the cohort of capsids from the same family, while TM-score measures the correspondence of CP structures. Notably, we also calculate the MDs between the model and its closely matched (reference) structure in evaluating how well the model compares with a known structure from the family. Moreover, the resulting models are relaxed by Amber energy minimization to relieve any steric clashes at the CP subunit interfaces in the predicted capsid model. Furthermore, we perform various capsid analysis - estimating BSAs at the unique subunit interfaces and corresponding association energies, accessible surface profiles, identifying the residue pairs that contact at these interfaces and simple parameters like diameters and net surface charge – that can also be compared with other members in the same virus family. The above metrics provide confidence measures of FoldaVirus predicted virus/capsid models. The predicated model coordinates can be downloaded for further analysis by the users.

Significantly, even when there is no structural information available directly for certain virus families, if the target CP sequence closely matches with the CPs in the structurally characterized families, FoldaVirus will be able to generate models for them. Going forward, in addition to sequence similarity as a measure to identify a type of capsid that a CP sequence is likely to form, we will implement the structural correspondence as another way to obtain such information, as the structural similarity is better conserved than the sequence similarity. In particular, the described approach overcomes the limitations of AlphaFold in generating the correct oligomers to represent IAUs, which is critical for building the accurate icosahedral capsids. Even though currently we have restricted the capsids that can be assembled to those with *T*-numbers less than or equal to 9, due to GPU memory limitations, the procedure can be readily expanded to generate more complex capsids exhibiting larger *T*-numbers (*T* > 9). Lastly, while we also locally implemented an equivalent workflow using AlphaFold3, which is 3-4 times faster than that uses AF2, we are unable to make it publicly available due to limitations on satisfying the terms and conditions of AF3 usage. However, the quality of resulting models from both the workflows appears to be very similar.

## Acknowledgements

This work was partially supported by the funds from Hormel Foundation to VSR.

## Author contributions

O.R.L., D.M.G. and V.S.R. conceptualized the study. O.R.L. and D.S.M.M. created all the workflows, Python scripts, web programming, generated the models and performed various tests. J.M. provided the guidance and assistance in building AlphaFold docker modules and setting up SLURM jobs on HPC cluster at the Hormel Institute. D.A.C. provided the guidance and assistance using Amber Tools and relaxation of atomic models. N.S.R., D.M.G. and V.S.R. supervised the study. O.R.L. and V.S.R. wrote and edited the manuscript with the feedback from all the authors.

---

**Supplementary Fig. 1.**
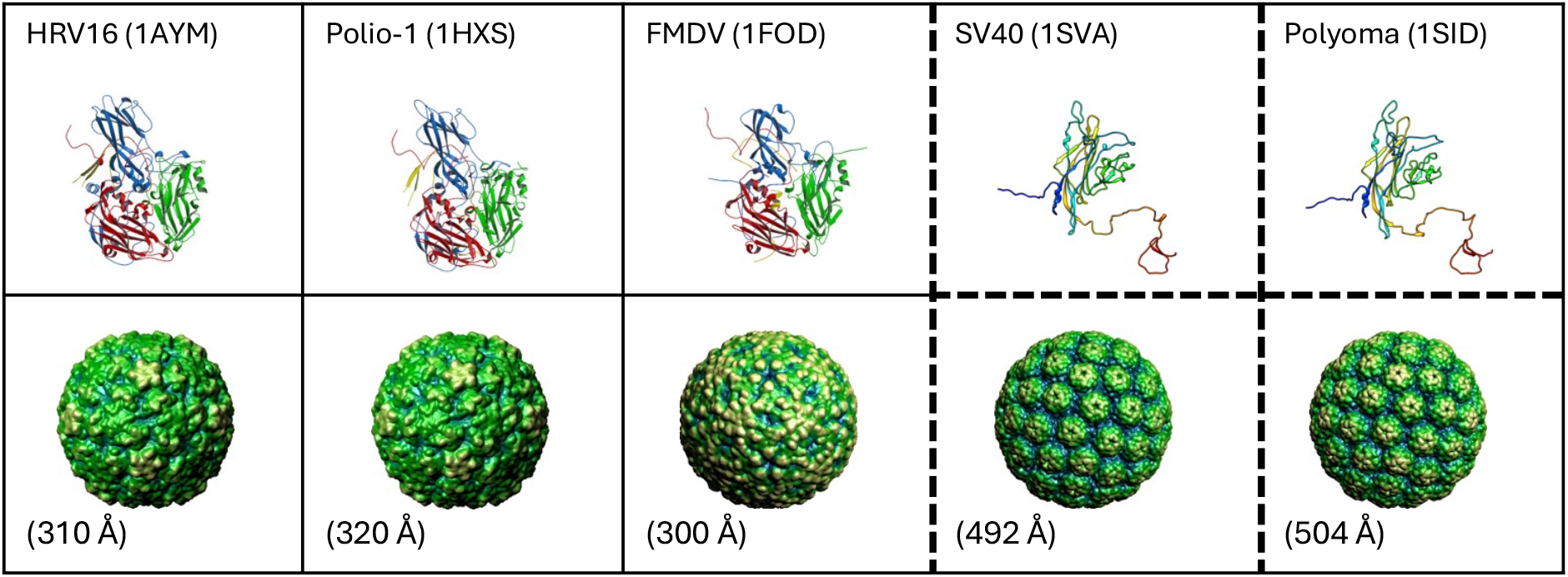
Ribbon diagrams representing the structures of CP protomers (top row) and corresponding rendered capsid surfaces (bottom row) of *Picornaviridae* and *Polyomaviridae* (shown in boxes with dotted lines) that form pseudo T=3 and T=7d capsids, respectively. Of note, different virus family pictures were not drawn on the same scale. The numbers in parentheses correspond to PDB-IDs (top row) and average diameters (bottom row) of the respective capsids.

**Supplementary Fig. 2.**
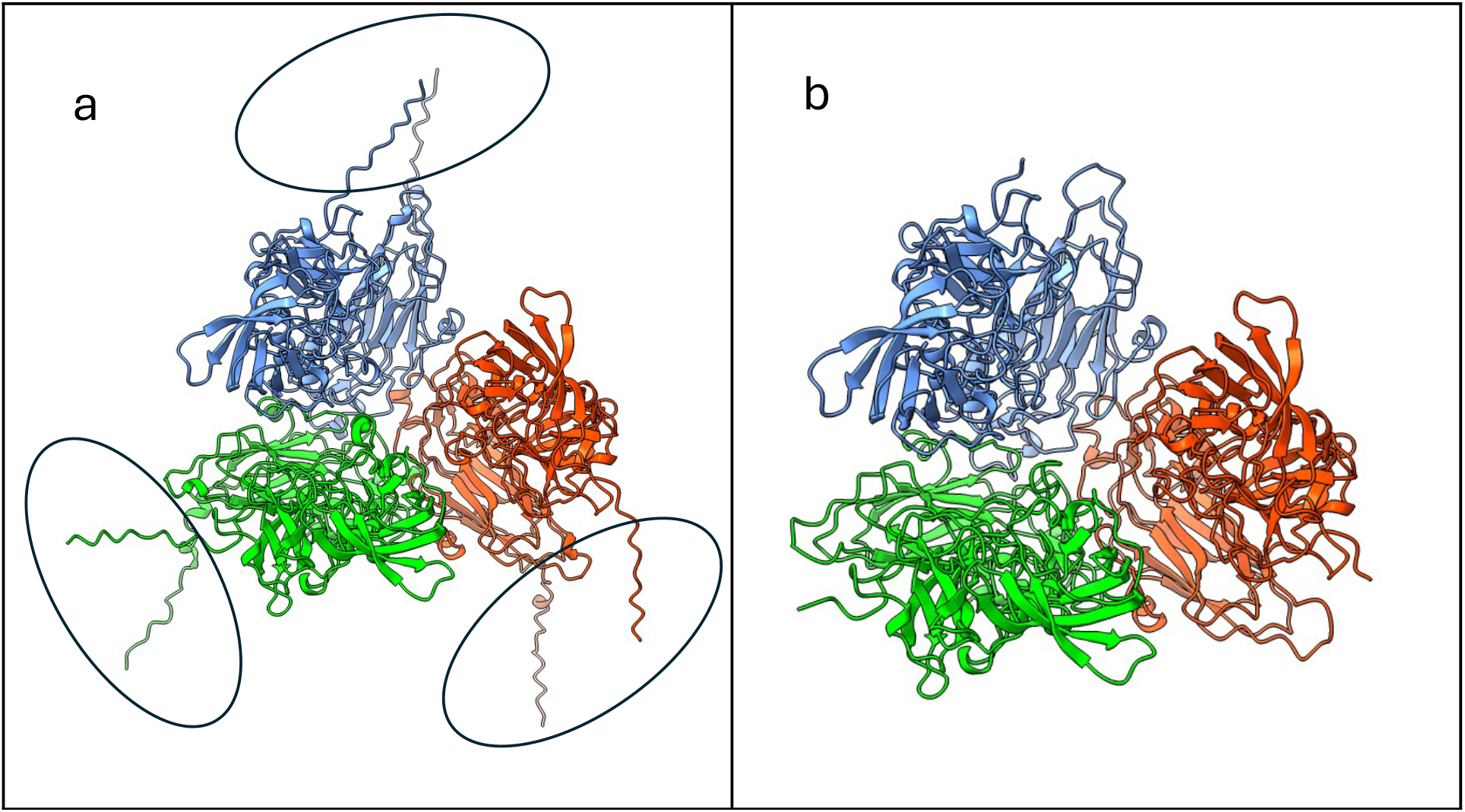
An example illustration of unstructured regions of the CP that were trimmed prior to generating the AF model. These regions often result in steric clashes while generating the complete capsids. a) AF-model of the icosahedral asymmetric unit (IAU) based on the full-length sequence of Norwalk virus CP (UniProt: Q83884). Structurally distinct subunits occupying the IAU are colored differently. The unstructured regions of the CPs located at the N and C-termini are encircled. b) AF-model of the IAU generated using the trimmed sequence of the above Norwalk virus CP shown in panel (a).

**Supplementary Fig. 3.**
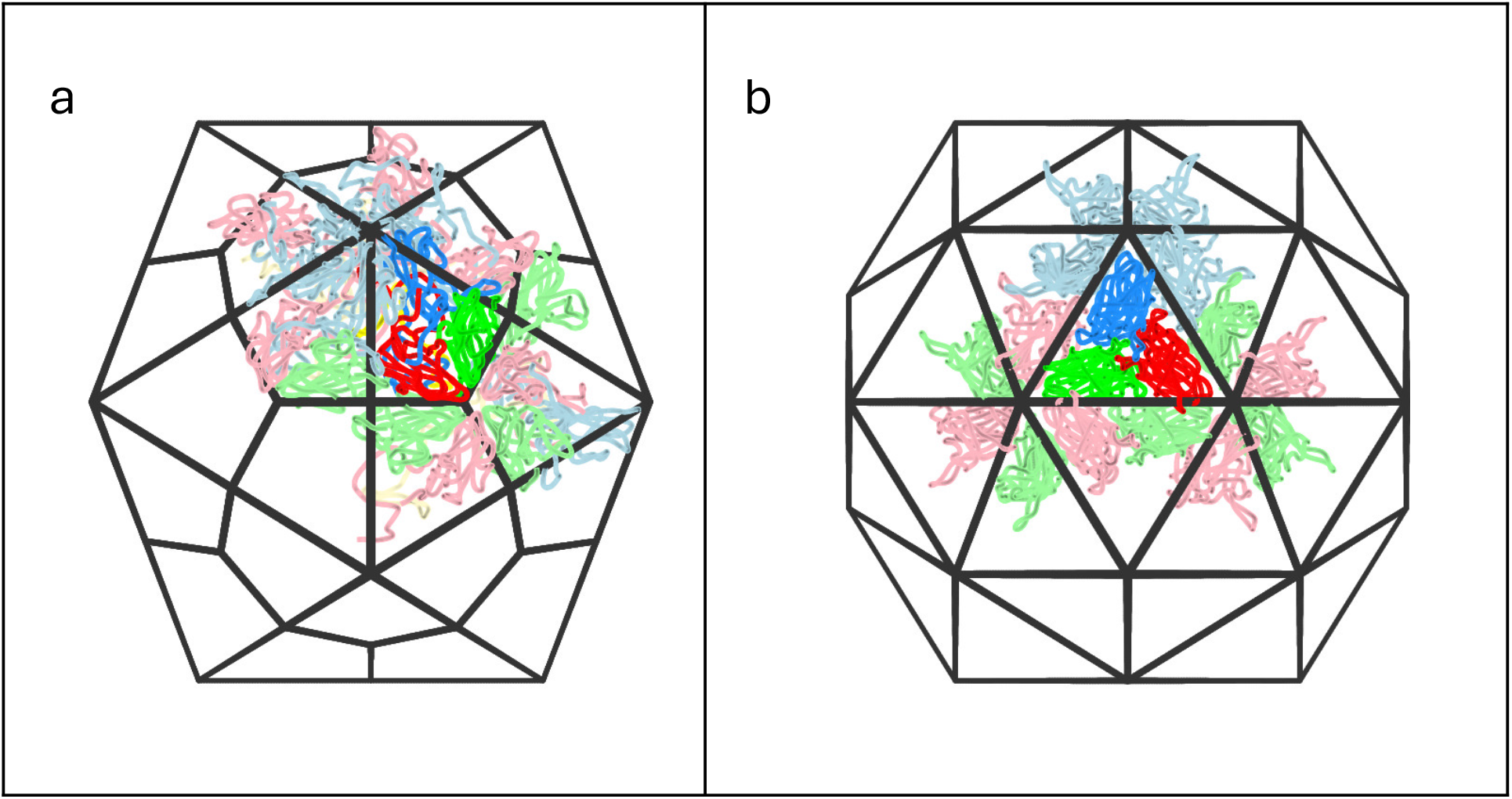
Schematic representations of the reference icosahedral asymmetric units (IAU) and the surrounding subunits used to generate partial capsids. **a)** A schematic of T=1 icosahedral lattice (black lines) with the reference IAU of a poliovirus (protomer) shown in dark colors, while the 15 surrounding subunits are shown in light colors which together represent the partial capsid. A similar scheme was used for all the capsids except those displaying T=3 icosahedral symmetry **b)** A schematic of T=3 icosahedral lattice (black lines) with the reference IAU of Black beetle virus shown in dark colors, while the 14 surrounding subunits are shown in light colors which together represent the partial capsid.

**Supplementary Fig. 4.**
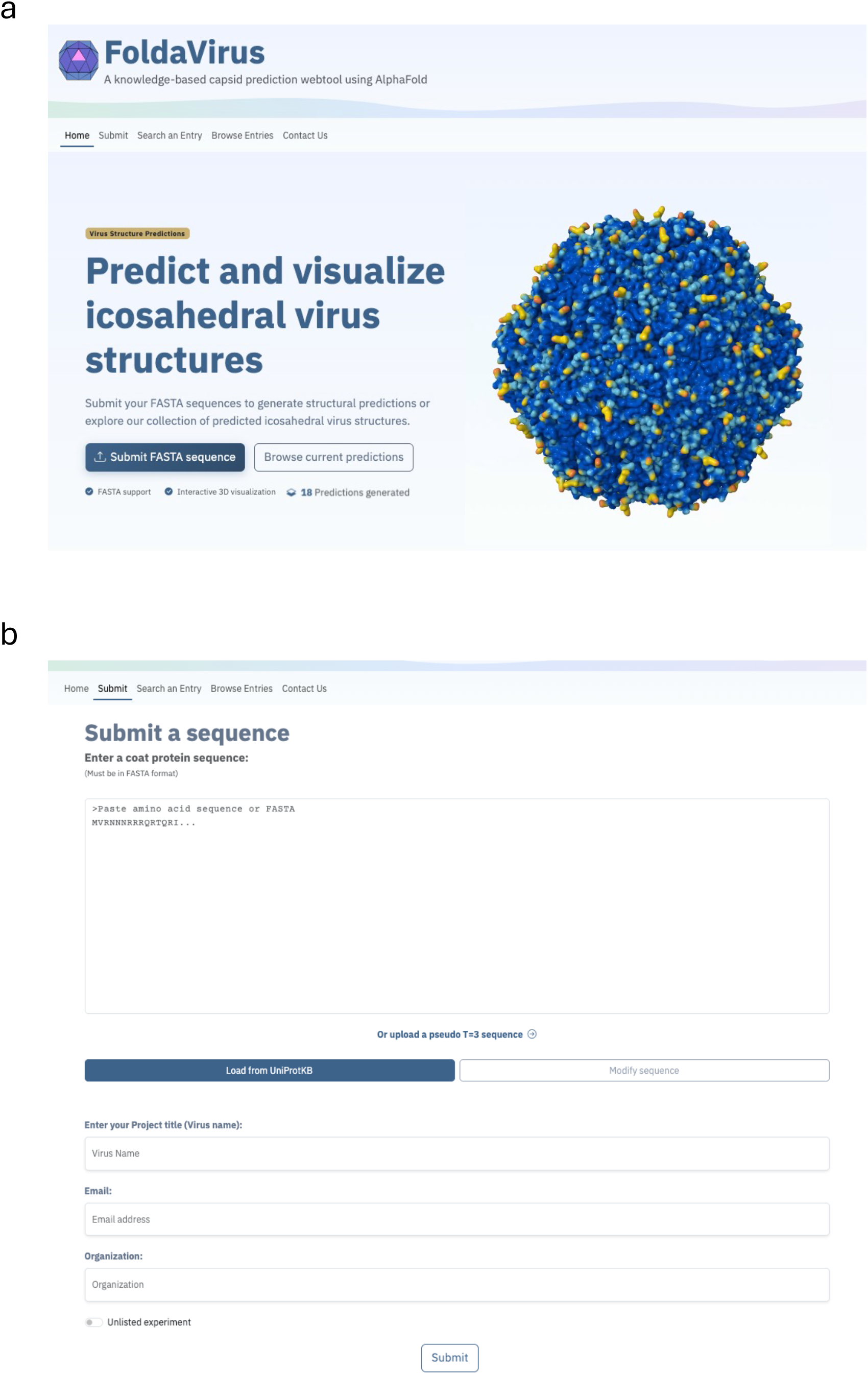
Web interface of FoldaVirus. **a)** Home page of Foldavirus (https://foldavirus.org) and **b)** web interface of user data submission page.

**Supplementary Fig. 5.**
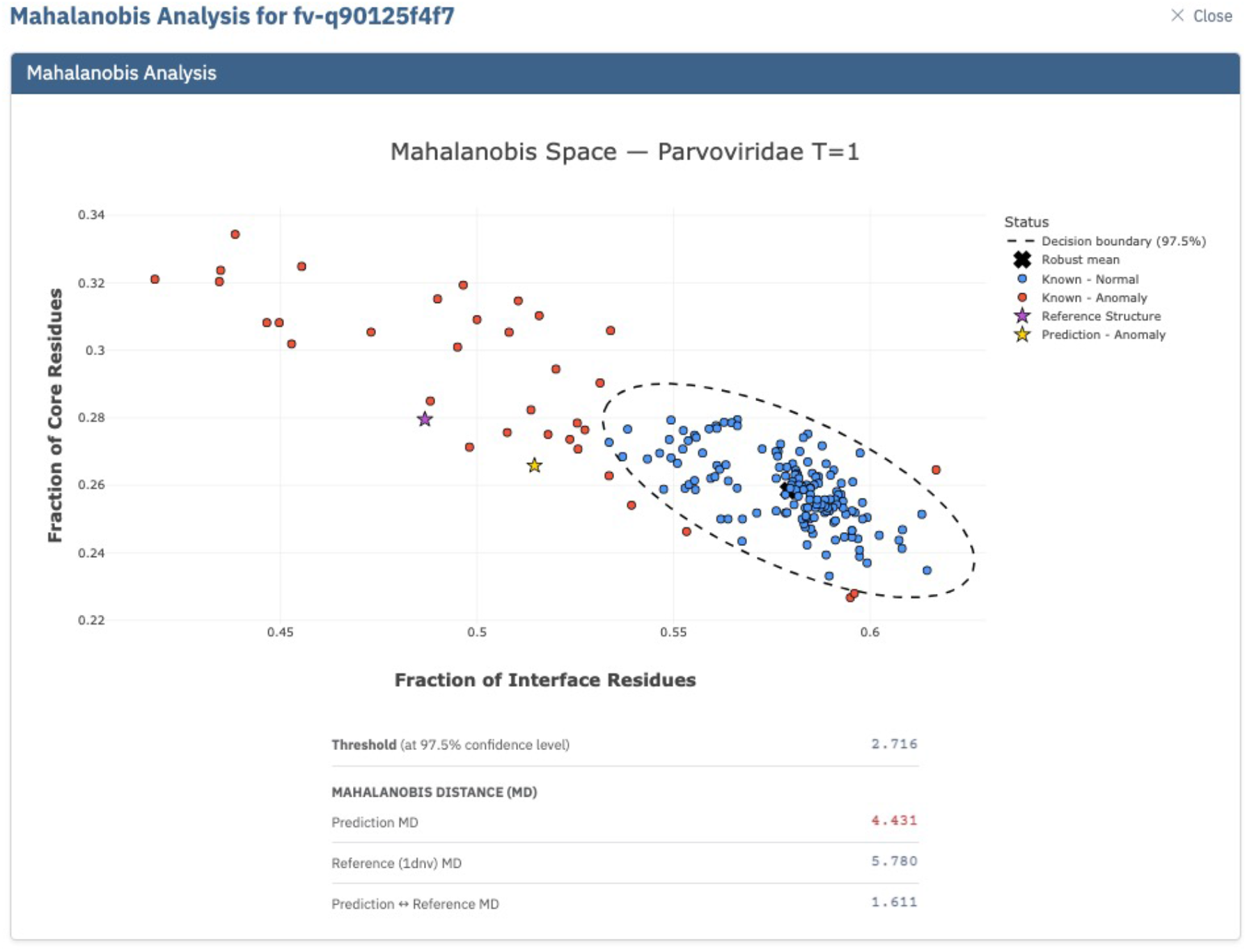
Mahalanobis plot of FoldaVirus model validation, a case study, where both the reference and predicted models are designated as outliers, but the model is closer to reference structure based on the pairwise Mahalanobis distance between them.

